# Transsynaptic neural circuit mapping of ventral hippocampus motivational control systems

**DOI:** 10.64898/2026.02.16.706218

**Authors:** Molly E. Klug, Mugil V. Shanmugam, Haoyang Huang, Don Arnold, Joel Hahn, Scott E. Kanoski

**Author notes:** Corresponding author: Scott E. Kanoski.

## Abstract

The ventral hippocampus contributes to food intake regulation and a range of motivational and memory processes, and its dysfunction is associated with several cognitive and behavioral disorders. However, its circuit-level organization remains incompletely understood. Ventral CA1 (CA1v) neurons send projections to several regions involved in motivational control. Here we focus on three major forebrain targets: the nucleus accumbens shell (ACBsh), medial prefrontal cortex (mPFC), and lateral hypothalamic area (LHA). We mapped the upstream and downstream circuitry of CA1v neurons defined by their projections to these target regions in rats using complementary transsynaptic anterograde and retrograde viral tracing approaches. Monosynaptic outputs to ACBsh, mPFC, and LHA were targeted using ATLAS, a novel transsynaptic anterograde viral approach that drives Cre recombinase in neurons receiving glutamatergic synaptic transmission from the CA1v. Second-order projections arising from these defined pathways were then mapped using a Cre-dependent anterograde viral tracing strategy. In parallel, upstream inputs to CA1v neurons projecting to each downstream target were mapped using a conditional retrograde glycoprotein-deleted rabies viral approach. Anterograde tracing revealed both shared and pathway-specific second-order targets, including bidirectional CA1v projections. Retrograde tracing confirmed expected inputs (e.g., CA3) and uncovered previously unrecognized cortical sources that differed across downstream projection-defined CA1v subpopulations. Together, these findings delineate pathway-specific, multinode circuits linking CA1v neurons to key motivational systems that may inform future therapeutic strategies for disorders involving ventral hippocampal dysfunction.

## Introduction

The hippocampus (HPC), most widely known for its role in learning and memory processes, plays an important role in the etiology of many neuropsychiatric disorders, including depression (Bagot et al., 2015; Medrihan et al., 2022), schizophrenia (Mikell et al., 2009; Tseng et al., 2009), and substance abuse disorders (Rogers and See, 2007; Bossert et al., 2016; Caban Rivera et al., 2023). The HPC also plays an important role in regulating reward-motivated behavior (Cooper et al., 2006; LeGates et al., 2018), influencing effort-based responding (Schmelzeis and Mittleman, 1996), behavioral inhibition (Abela et al., 2013; Assari et al., 2025), and impulsivity (Abela et al., 2013; Hsu et al., 2018; Noble et al., 2019; Masuda et al., 2020). Recent and emerging work also reveals a critical role for the HPC in regulating appetite, caloric consumption, and body weight regulation (Kanoski and Grill, 2017; Stevenson and Francis, 2017; Barbosa et al., 2023), in part by modulating satiation and meal size control (Suarez et al., 2020), promoting cue-potentiated eating (Kanoski et al., 2013), and encoding meal-related memories (Hannapel et al., 2017, 2019; Décarie-Spain et al., 2022, 2025a; Yang et al., 2025). A better understanding of the neural circuitry via which the HPC contributes to the etiology of obesity, neuropsychiatric disorders, and disordered motivational outcomes could help to identify novel therapeutic targets.

While the hippocampus is often referred to a singular structure, this brain region contains several distinct subregions, each with their own distinguishable cytoarchitecture (Andersen, 2007), genetic signature (Bienkowski et al., 2018), and connectivity (Kanoski and Grill, 2017). In addition to the CA fields (1, 2, and 3) and the dentate gyrus (DG) subregional cellular distinctions that extend across the longitudinal axis, the rodent hippocampus is often divided into two larger subregions: the anterior “dorsal” hippocampus (dHPC) and the posterior “ventral” (vHPC), with some distinctions also including an “intermediate” HPC region (Fanselow and Dong, 2010).

The dHPC and vHPC subregions in the rodent are analogous to the posterior and anterior portions of the HPC in humans, respectively (Lee et al., 2019). The term ‘hippocampal formation’ refers to the hippocampus proper (CA and DG fields), entorhinal cortex (EC), and subiculum (SUB) (O’Keefe and Nadel, 1978). Both the dorsal and ventral hippocampal regions have been shown to play a critical role in food intake and food-motivated behavior (Kanoski and Grill, 2017; Hannapel et al., 2019), and these two subregions share common downstream targets – including the nucleus accumbens, subiculum, and septal nuclei (Fanselow and Dong, 2010; Trouche et al., 2019; Barnstedt et al., 2024; Ibrahim et al., 2024). Despite these common downstream targets, these two hippocampal subregions are largely considered to be functionally and anatomically distinct (Fanselow and Dong, 2010), with the vHPC established as being particularly important in food intake and food-motivated behaviors. More specifically, work from our group and others has established the ventral CA1 (CA1v) as an important regulator of food intake control, both with regards to cue-stimulated appetite, and satiation control (Hsu et al., 2015b, 2015a, 2018; Suarez et al., 2020; Décarie-Spain et al., 2025b).

While CA1v is a critical node in the classic ‘tri-synaptic circuit’ of the hippocampus via its inputs from CA3v and via subiculum outputs (Andersen, 2007), several important extra-hippocampal formation projections have been established. Our group and others have shown that the CA1v sends projections to the lateral septum (LS) (Sweeney and Yang, 2015; Kosugi et al., 2021; Décarie-Spain et al., 2022), the medial prefrontal cortex (mPFC) (Chudasama et al., 2012; Hsu et al., 2018), the nucleus accumbens shell (ACBsh) (Bagot et al., 2015; Yang et al., 2020; Zhou et al., 2020; Tsai et al., 2022; Klug et al., 2025; Patterson et al., 2025), the basolateral amygdala (BLA) (Jimenez et al., 2018), and the lateral hypothalamic area (LHA) (Cenquizca and Swanson, 2006; Hahn and Swanson, 2010, 2012; Hsu et al., 2015c; Suarez et al., 2020; Décarie-Spain et al., 2025b) . Downstream projections from the CA1v to most of these targets have been shown to be relevant to food-motivated behavior and/or food consumption (Hsu et al., 2018; Décarie-Spain et al., 2022, 2025b), thus making them important potential targets for future obesity therapeutics.

Recent advances in viral-based neural tract tracing allow for transsynaptic-level insights, thus offering the potential to move beyond simple monosynaptic connections between regions to identify complex multi-node neuronal circuitry. To gain such insights for three distinct CA1v projection target pathways that are functionally linked with the control of food intake control, here we use anterograde and retrograde transsynaptic viral tracing approaches to identify both downstream and upstream connections from and to, respectively, the CA1v-mPFC, CA1v-ACBsh, and CA1v-LHA projection pathways. The novel transsynaptic ATLAS approach (Rivera et al., 2025) provides us with the unique opportunity to probe this circuitry further by establishing what brain regions lie downstream of glutamatergic synaptic transmission from CA1v-mPFC, CA1v-ACBsh, and CA1v-LHA projections. This is in contrast to previous anterograde transsynaptic tracing approaches which may present synaptic leakage or retrograde transmission that are distinct from physiological glutamate transmission (Zingg et al., 2017). Further, given that all extrinsic projections from the CA1v are glutamatergic, the ATLAS approach represents a comprehensive tool to transsynaptically trace from these neurons. Additionally, we utilized a transsynaptic glycoprotein-deleted retrograde rabies virus approach (Wall et al., 2010; Callaway and Luo, 2015) to identify inputs into CA1v neurons based on their specific projection targets to mPFC, ACBsh, or LHA. Thus, in this paper we elucidate three distinct multi-node neuronal circuits centering on CA1v projections that are functionally relevant to food intake and body weight regulation (Figure 1A).

**Figure 1.**
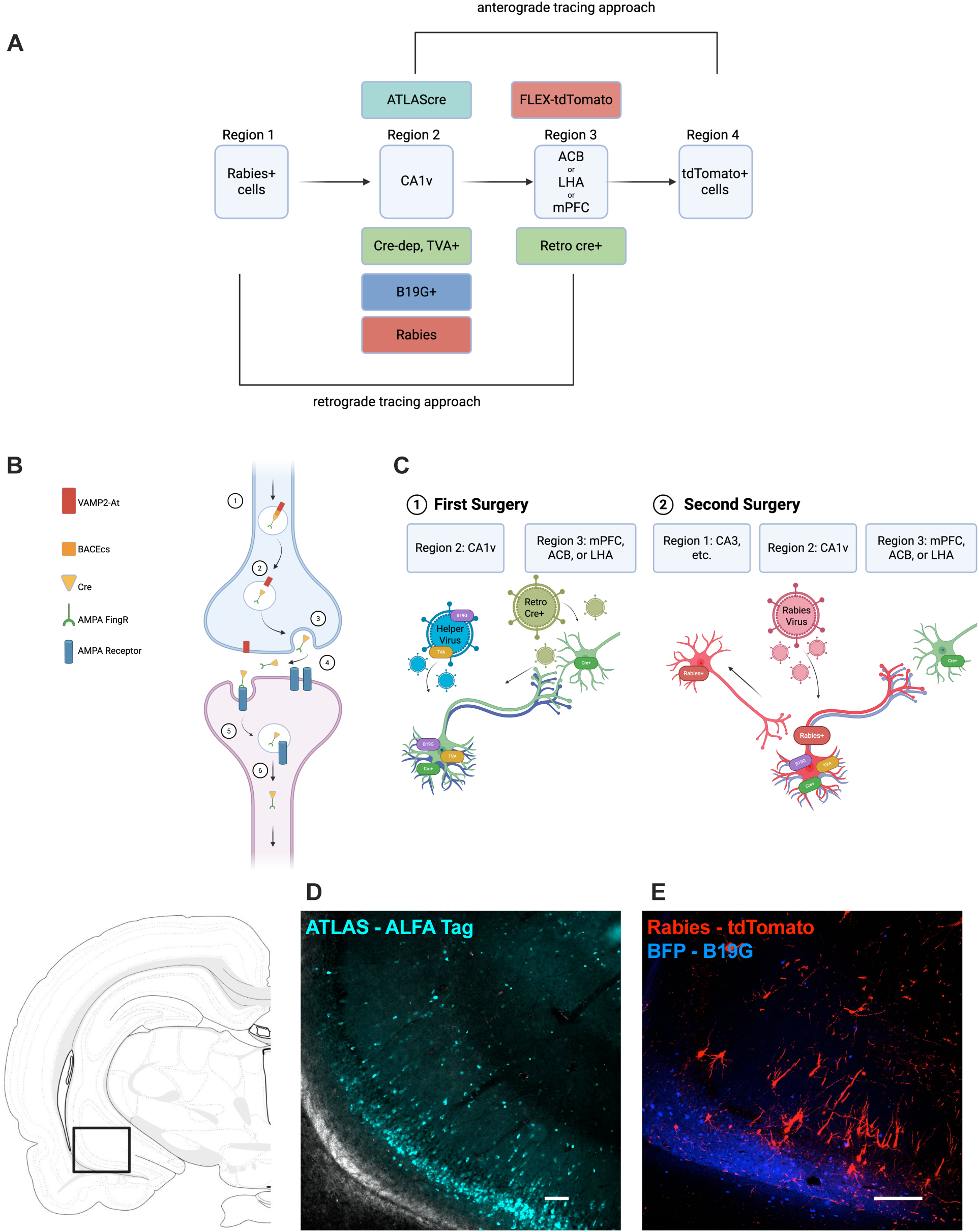
Methods for multi-node, multi-synaptic pathway tracing. **A.** Schematic depicting both anterograde and retrograde multi-synaptic tracing approach. **B.** Schematic depicting mechanism of action for ATLASsncre virus; 1) ATLASsncre is expressed in the presynaptic starter cell and targeted to the synapse via a VAMP2 domain. 2) A BACEcs is cleaved, 3) allowing AF-Cre to enter the synaptic cleft. 4) AF binds to GluA1 on the postsynaptic cell and 5) is endocytosed. 6) AF-cre moves to the nucleus of the postsynaptic cell, driving Cre expression (adapted from (Rivera et al., 2025). **C.** Schematic depicting surgical approach for pseudo-rabies viral tracing approach; 1) in the first surgery, the rabies helper virus is infused into the CA1v driving expression of TVA receptor and rabies G-glycoprotein (B19G; necessary for transneuronal transfer of the rabies virus), along with a retro-cre inducing virus in 1 of 3 possible downstream regions. 2) A week later, in a second surgery, the rabies virus is infused into the CA1v, driving rabies expression in CA1v neurons which project to 1 of 3 possible downstream regions via EnvA expression (the ligand for the TVA receptor). **D.** ATLASsncre **E.** Representative image of Rabies and Helper viruses in the CA1v (Scale bar, 100um).

## Methods and Materials

### Animals

Male Sprague-Dawley rats (Envigo, Indianapolis, IN; postnatal day [PND] 60-70; 250-275g on avg upon arrival) were individually housed in a temperature-controlled vivarium with *ad libitum* access (except where noted) to water and food (LabDiet 5001, LabDiet, St. Louis, MO) on a 12h:12h reverse light/dark cycle. All procedures were approved by the Institute of Animal Care and Use Committee at the University of Southern California.

### Surgical procedures

For all surgical procedures, rats were anesthetized and sedated via intramuscular injections of ketamine (90 mg/kg), xylazine (2.8 mg/kg), and acepromazine (0.72 mg/kg). Animals were shaved, surgical site was prepped with iodine and ethanol swabs, and animals were placed in a stereotaxic apparatus for stereotaxic injections. Viruses were delivered using a microinfusion pump (Chemyx, Stafford, TX, USA) connected to a 33-gauge microsyringe injector attached to a PE20 catheter and Hamilton syringe. Flow rate was calibrated and set to 5 µl/min. Injectors were left in place for 2 min post injection, and animals were sutured following injections. Rats were also given analgesic (subcutaneous injection of ketoprofen [5mg/kg]) after surgery and once daily for 3 subsequent days thereafter. All rats recovered for at least three-weeks post-surgery prior to tissue collection to allow for viral transduction and expression.

### Viral injections

For neural tracing from second-order neurons receiving projections from the defined CA1v monosynaptic projection pathways, AAV8-ATLASsncre and pAAV-DIO0BACE-HA (ATLASsncre; Gift from the Arnold Lab (Rivera et al., 2025); available on Addgene, ID 232351 and ID232353; 200nL per side) was unilaterally injected at the following coordinates: −4.9 mm anterior/posterior (AP), +-4.8 mm medial/lateral (ML), −7.8 mm dorsal/ventral (DV) (0 reference point at bregma for ML, AP, 0 reference point at skull surface near injection site for DV). Additionally, an AAV1-CAG-Flex-tdTomato-WPRE-bGH, (FLEX-tdTomato; UPenn Vector Core; 200nL per side) was unilaterally injected at one of the following coordinates, to allow for anterograde neural tracing from CA1v targets:

ACBsh: +1.2 mm AP, +-1.0 mm ML, −6.75 mm DV (0 reference point at bregma for ML, AP, 0 reference point at dura for DV).

LHA: -2.9 mm AP, +-1.0 mm and +- 1.6mm ML, −8.8 mm DV (0 reference point at bregma for all coordinates; skull surface at bregma for DV).

mPFC: 2.5 mm AP, +-0.5 mm and +- 1.6mm ML, −4.0 mm DV (0 reference point at bregma for all coordinates).

For retrograde tracing from from CA1v neurons based on their specific monosynaptic projection targets, surgeries were performed as previously described (Liu et al., 2017). The helper viruses pAAV-syn-FLEX-splitTVA-EGFP-tTA and pAAV-TREtight-mTagBFP2-B19G (Helper virus; Addgene, ID100798 and ID100799) were mixed 1:1 and unilaterally injected at the CA1v (600nL; see coordinates above), and a retrograde cre-expressing virus AAV-eSYN-EGFP-T2A-iCre-WPRE (Retro cre; Vector BioLabs, Cat #VB4855) was injected into either the ACB, LHA, or mPFC (400nL; see coordinates above). Seven days later, the rabies virus EnvA-SAD-B19-RVΔG tdTomato (Rabies; UC Irvine, Center for Neural Circuit Mapping, Cat. No: EnvA-RV-2, California, USA) was unilaterally injected into the CA1v (400nL), allowing retrograde tracing from CA1v neurons that project to either the ACBsh, LHA, or mPFC. Seven days later (14 days after the initial surgery), animals were euthanized, and brains were collected as described below.

### Immunohistochemistry (IHC)

Rats were anesthetized and sedated with a ketamine (90 mg/kg)/xylazine (2.8 mg/kg)/acepromazine (0.72 mg/kg) cocktail, then transcardially perfused with 0.9% sterile saline (pH 7.4) followed by 4% paraformaldehyde (PFA) in 0.1 M borate buffer (pH 9.5; PFA). Brains were dissected out and post-fixed in PFA with 15% sucrose for 24 h, then flash frozen in isopentane cooled in dry ice. Brains were sectioned to 30-µm thickness on a freezing microtome. Sections were collected in 5 series and stored in antifreeze solution at −20 °C until further processing.

Successful virally-mediated transduction was confirmed postmortem in all animals via IHC staining using immunofluorescence-based antibody amplification to enhance the fluorescence. The following antibodies and dilutions were used: rabbit anti-RFP (1:2000, Rockland Inc., Limerick, PA, USA), and chicken anti-GFP (1:1000, Cat. no. ab13970, Abcam, Cambridge, UK), ALFA-TAG (1:1000, Cat No: N1502, NanoTag Biotechnologies, Gottingen, Germany). Antibodies were prepared in 0.02 M potassium phosphate-buffered saline (KPBS) solution containing 0.2% bovine serum albumin and 0.3% Triton X-100 at 4 °C overnight. After thorough washing with 0.02 M KPBS, sections were incubated in secondary antibody solution. All secondary antibodies were obtained from Jackson Immunoresearch and used at 1:500 dilution at 4 °C, with overnight incubations (Jackson Immunoresearch; West Grove, PA, USA). Sections were mounted and coverslipped using 50% glycerol in 0.02 M KPBS and the edges were sealed with clear nail polish. Photomicrographs were acquired using either a Nikon 80i (Nikon DS-QI1,1280X1024 resolution, 1.45 megapixel) under epifluorescence or darkfield illumination.

### Axonal Density and Cell Count Quantification

Data were entered using a custom built data-entry platform (Axiome C, created by JDH) built around Microsoft Excel software and designed to facilitate entry of data points for all gray matter regions across their atlas levels as described in a rat brain reference atlas: Brain Maps 4.0 (Swanson, 2018). The Axiome C approach was used previously to facilitate the analysis of downstream projection targets in pathway tracing studies (Décarie-Spain et al., 2022; Hahn et al., 2022). For anterograde labeling, an ordinal scale, ranging from 0 (absent) to 7 (very strong) was used to record the qualitative weight of axonal density. For retrograde labeling, RFP+ cells were quantified using ImageJ (Rueden et al., 2017). An average, total, and most common value was then obtained for each region across its atlas levels for which data were available. These data are summarized graphically for a representative animal on a brain flatmap summary diagram generated using a software-based approach (Hahn and Duckworth, 2023; and partially adapted from Décarie-Spain et al., 2022), as well as on a bar graph with collapsing across some subregions (e.g., for LHA and LS) for the latter data depiction.

### Experimental Design and Statistical Analyses

Data are represented as either modes, means, maximums, or total values; data analysis details can be found in the figure legends.

## Results

### Anterograde tracing of second-order projections from three distinct brain regions receiving monosynaptic projections from the CA1v

In order to characterize downstream brain regions of the CA1v to mPFC, CA1v to ACBsh, and CA1v to LHA pathways, we employed a novel transsynaptic viral tracing approach, ATLAS, to label downstream neurons (and their projections) in a strictly anterograde manner mediated by physiological glutamatergic synaptic transmission (Rivera et al., 2025) (Figure 1B). The ATLASsncre AAV drives expression of the AMPA.FingR (AF; FingR stands for fibronectin intrabody generated with mRNA display) attached to a synaptobrevin 2 (VAMP2) domain fused to a Synaptotagmin nanobody, thus targeting the ATLAS virus to the synapse. Additionally, the virus contains a beta secretase (BACE) cleavage site (BACEcs) between the AF and the VAMP2 domain, allowing endogenous BACE to cleave the AF from the VAMP2 domain and exit the lumen of the synaptic vesicle. Once in the synaptic cleft, AF, and its Cre payload, can bind to GluA1 on the postsynaptic cell, thereby being endocytosed and allowing for transmission of Cre in the postsynaptic cell (Figure 1B). We unilaterally injected ATLASsncre in the CA1v (Figure 1D) and FLEX-tdTomato in either the mPFC, ACBsh, or LHA to trace downstream targets of these pathways (Figure 1A).

Forebrain- and midbrain-wide neuroanatomical imaging and quantification of axon density revealed that mPFC neurons receiving projections from the CA1v send dense projections throughout the brain, with the medial septum (MS) and CA1v having among the highest most common value in terms of axon density (Figure 2A-C), while the CA1v and periventricular hypothalamic nucleus (PVp) had high average axon density (Supp. Figure 1A,B), and the lateral septum (LS), lateral hypothalamic area (LHA), and the anterior cingulate area (ACA) showed among the greatest total axon density (Supp. Figure 2A,B) on the side ipsilateral to the injection site. On the contralateral side, the LS and olfactory tubercle (OT) had among the highest mode for axon density, while the septofimbrial nucleus (SF) and anteromedial thalamic nucleus (AM), and LS and paraventricular thalamic nucleus (PVT), scored the highest in terms of average and total axon density respectively (Supp. Figure 1A,B; Supp. Figure 2A,B). When analyzing for the maximum axon density, the RE scored among the highest on both the ipsilateral and contralateral side of the CA1v-mPFC pathway (Supp. Figure 3A,B).

**Figure 2.**
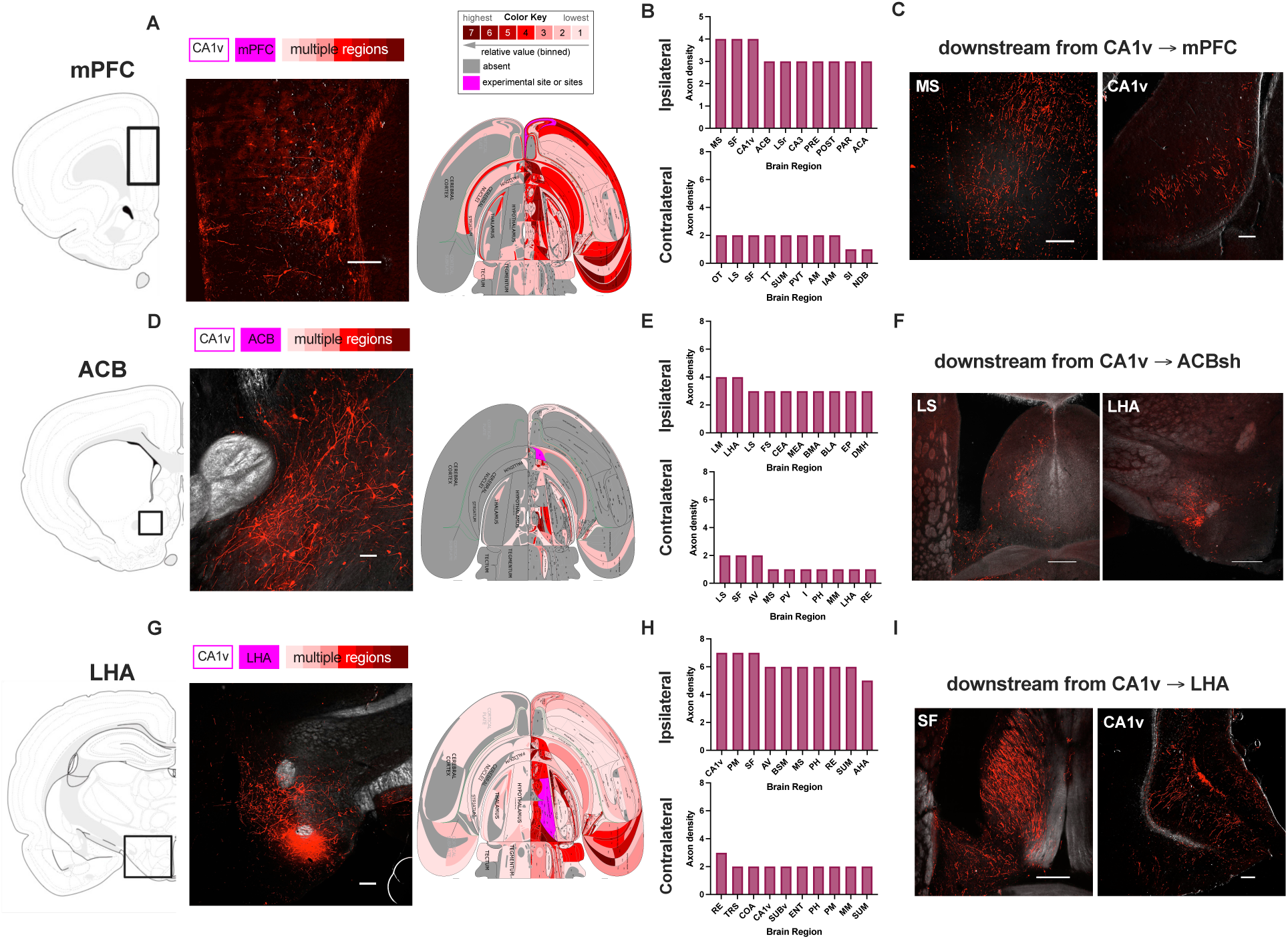
Most common axon density in anterograde tracing of 2nd-order projections from CA1v pathways. **A.** Representative image (left) and flatmap (right) depicting most common axon density for CA1v-mPFC anterograde tracing. **B**. Ipsilateral (top) and contralateral (bottom) graphs depicting most common axon density rating in CA1v-mPFC pathway. **C.** Representative images depicting axon density in the RE (left) and CA1v (right). **D.** Representative image (left) and flatmap (right) depicting most common axon density for CA1v-ACB anterograde tracing. **E**. Ipsilateral (top) and contralateral (bottom) graphs depicting most common axon density rating in CA1v-ACB pathway. **F.** Representative images depicting axon density in the LS (left) and LHA (right). **G.** Representative image (left) and flatmap (right) depicting most common axon density for CA1v-LHA anterograde tracing. **H**. Ipsilateral (top) and contralateral (bottom) graphs depicting most common axon density rating in CA1v-LHA pathway. **I.** Representative images depicting axon density in the FS (left) and CA1v (right).

The brain regions with among the highest mode of axon density that are innervated by the Ca1v-ACBsh pathway include the LS and the LHA (Figure 2D-F), while the innominate substance (SI), lateral mamillary nucleus (LM), and mediodorsal thalamic nucleus medial part (MDm) had among the greatest average axon density on the ipsilateral side (Supp. Figure 1C,D). When examining projections contralateral to the injection site, the LS and the SF were among the highest scoring regions for both mode and average of axon density (Figure 2E; Supp. Figure 1C,D) Finally, the LHA, bed nuclei of terminal stria (BST), and the LS had particularly high total axon density in the CA1v-ACBsh pathway on the ipsilateral side, and the LS and PVT had among the highest total axon density on the contralateral side (Supp. Figure 2C,D). The LS also had strong maximum axon density score on both the ipsilateral and contralateral sides (Supp. Figure 3C,D).

LHA neurons receiving projections from the CA1v project to multiple downstream brain regions, with the bed nucleus of medullary stria (BSM) and CA1v being among the most robust downstream targets when analyzing by most common axon density value (Figure 2G-I) and by average axon density value on the ipsilateral side (Supp. Figure 1E,F). On the side of the brain contralateral to tracing injections, the RE and triangular septal nucleus (TRS) had among the highest mode for axon density, while the RE and medial mammillary nucleus (MM) had the highest average axon density (Figure 2H; Supp Figure1E,F). When analyzing for total axon density per region the BST, LS, and periaqueductal gray (PAG) scored among the highest on the ipsilateral side, and the LS and PAG scored among the highest in terms of contralateral projections (Supp. Figure 2E,F). Finally, when analyzing for maximum axon density in second-order projections, the CA1v and dorsomedial hypothalamus (DMH) scored among highest on the ipsilateral side, while the mammillary nucleus (MM) and the medial septum (MS) scored among highest on the contralateral side (Supp. Figure 3E,F).

### Retrograde tracing from CA1v neurons that project to either the ACBsh, LHA, or mPFC

To characterize the upstream brain regions of the CA1v-ACBsh, CA1v-LHA, and CA1v-mPFC pathways, we utilized a monosynaptic retrograde rabies viral approach. A helper virus, encoding for both a cre-dependent TVA receptor and the rabies G glycoprotein, were injected into the CA1v, along with a retro-cre virus injection in either the mPFC, ACBsh, or LHA. Rabies G is a necessary glycoprotein essential for transneuronal transfer of the rabies virus, while the TVA receptor is the receptor that the envelope protein from avian ASLV type A (EnvA) uses for cell entry. Importantly, TVA is not expressed in mammalian neurons, meaning that this combination of viral infusions ensures that the rabies G protein is expressed in all CA1v cells, while the TVA receptor expression is only driven in CA1v cells that project to one of our downstream regions of interest (ROI) (Figure 1C). In a second surgery, a pseudotyped rabies virus is infused in the CA1v, one which lacks the Rabies G protein but expresses EnvA (Figure 1C). Collectively, this surgical approach allows us to perform monosynaptic and retrograde tracing solely from CA1v neurons which further project to either the mPFC, ACB, or LHA (Figure 1A).

Our results reveal that the CA3 and the ventral subiculum (SUBv) give robust input to both the CA1v-mPFC and CA1v-LHA pathways (Figure 3A-C, G-I). While the CA1v-ACBsh also receives input from the CA3, a majority of inputs to this pathway are cortical, including brain regions such as the posterior amygdalar nucleus (PA) and the cortical nucleus of the amydgala (COA) (Figure 3D-F). Additionally, the postpiriform transition area (TR) and the basolateral amygdala (BLA) are among the top projections to both the CA1v-mPFC and CA1v-ACBsh pathways, while each projects minimally to the CA1v-LHA pathway (Figure 3A-F). Of note, the VTA also sends projections to the CA1v-mPFC pathway, albeit minimally (Figure 3A,B).

**Figure 3.**
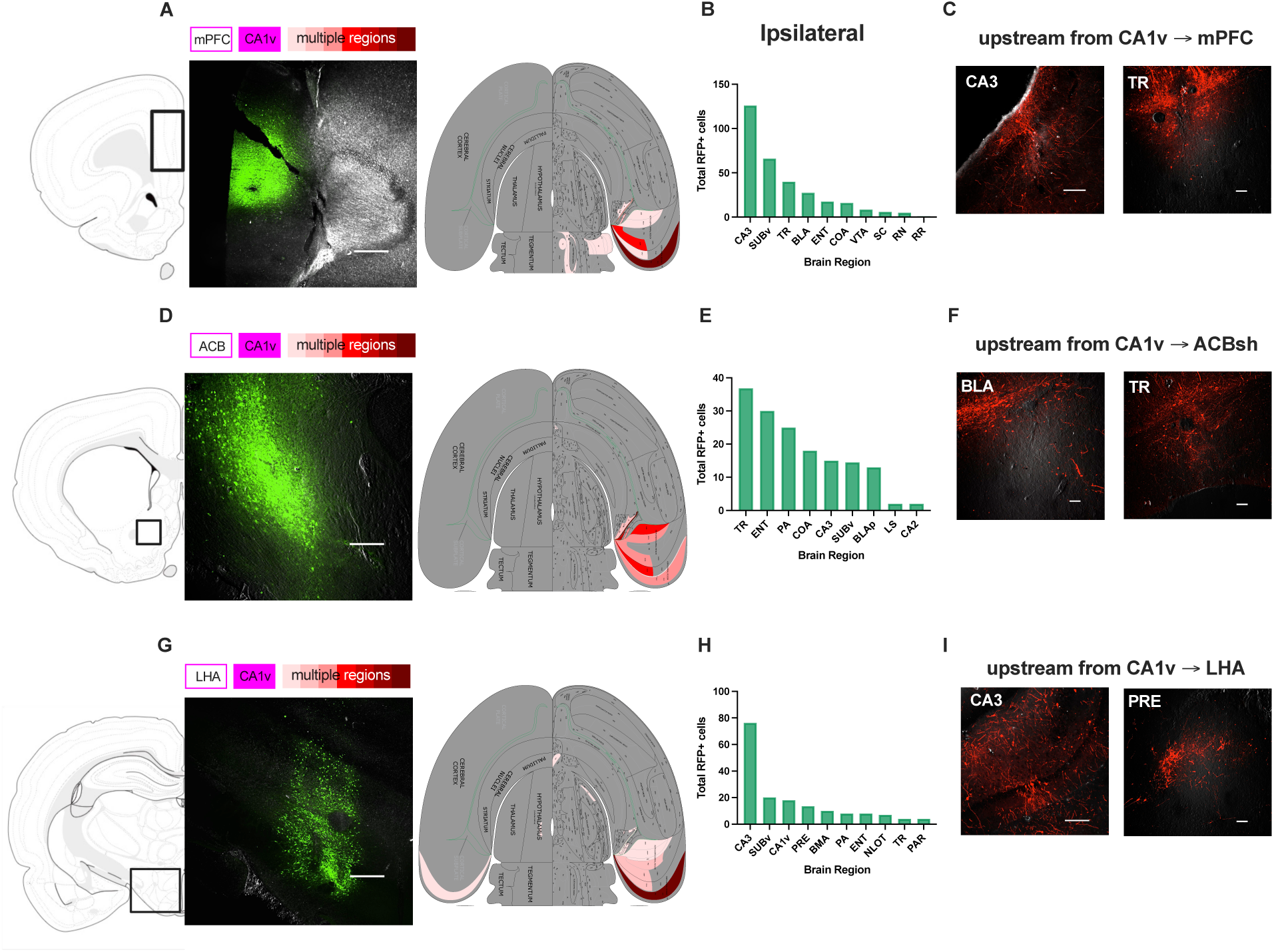
Total RFP+ cells in retrograde tracing of primary inputs to CA1v pathways. **A.** Representative image (left) and flatmap (right) depicting total RFP+ cell bodies for CA1v-mPFC retrograde tracing. **B**. Ipsilateral graph depicting total RFP+ cells for retrograde tracing in CA1v-mPFC pathway. **C.** Representative images depicting cell bodies in the TR (left)and CA3v (right). **D.** Representative image (left) and flatmap (right) depicting total RFP+ cell bodies for CA1v-ACB retrograde tracing. **E**. Ipsilateral graph depicting total RFP+ cells for retrograde tracing in CA1v-ACB pathway. **F.** Representative images depicting cell bodies in the BLA (left) and TR (right)**G.** Representative image (left) and flatmap (right) depicting total RFP+ cell bodies for CA1v-LHA retrograde tracing. **H**. Ipsilateral graph depicting total RFP+ cells for retrograde tracing in CA1v-LHA pathway. **I.** Representative images depicting cell bodies in the CA3v (left) and PRE (right).

## Discussion

The ventral hippocampus, particularly the CA1v subregion, has recently been implicated as a critical player in the regulation of appetite, food intake, and body weight regulation (Kanoski and Grill, 2017; Hsu et al., 2018; Noble et al., 2019). Further, perturbations in ventral hippocampus signaling is associated with various motivational and psychiatric deficits (Schmelzeis and Mittleman, 1996; Tseng et al., 2009; Bagot et al., 2015; Zhou et al., 2019; Muir et al., 2020; Medrihan et al., 2022). A better understanding of CA1v pathways regulating motivation for food and other reinforcers could facilitate the development of novel therapeutics for the treatment of obesity and neuropsychiatric disorders. Recent advances in transsynaptic anterograde tracing technology (Rivera et al., 2025), paired with established transsynaptic retrograde tracing approaches, have allowed us to uncover multi-node neuronal circuits connecting the CA1v to brain regions crucial for reward, emotional, and contextual processing. Our results outline three distinct, 4-node, multi-synaptic pathways anchored from known monosynaptic CA1v projection pathways previously implicated in motivational control. Using a unique combination of tracing approaches, we reveal both upstream and second-order downstream targets of the CA1v-mPFC, CA1v-ACBsh, and CA1v-LHA projection pathways. It has been well established that the CA1v-mPFC (Phillips et al., 2019), CA1v-ACB (Caban Rivera et al., 2023), and CA1v-LHA (Décarie-Spain et al., 2025b) projections play a role in hippocampal-dependent memory, but there is also a clear role for these pathways in both food intake and food-motivated behaviors (Chudasama et al., 2012; Hsu et al., 2015b, 2018; Trouche et al., 2019; Suarez et al., 2020; Klug et al., 2025).

One notable outcome of this study is that there are several common downstream brain regions receiving projections from these distinct CA1v-derived pathways. For example, the LS, known for its role in motivation and emotion (Wirtshafter and Wilson, 2021), is not only a monosynaptic target of CA1v neurons (Décarie-Spain et al., 2022), but also emerged as a common robust second-order target of both the CA1v-mPFC and CA1v-ACBsh pathways. In this capacity, the LS may receive integrated multi-modal information filtered through the HPC to mediate goal-directed behavior. Additionally, the LS is increasingly being recognized for its role in eating-related behavior (Sweeney and Yang, 2015; Kosugi et al., 2021), and previous research identified a role for melanin-concentrating hormone (MCH) signaling in the LS in promoting food intake (Payant et al., 2025). Given that MCH-producing neurons in the LHA that project to the CA1v have also been implicated in food-motivated behavior (Noble et al., 2019), these results raise the possibility that the LS is a critical hub integrating energy balance-relevant information from hippocampal-originating cortical and striatal pathways, as well as directly from hypothalamic MCH signaling.

Another shared downstream target of these pathways is the BST. Both the CA1v-ACBsh and CA1v-LHA pathways send robust 2nd-order projections to the BST, a brain region heavily implicated in the integration of motivational, affective, and stress-related signals (Wang et al., 2025). Interestingly, the BST has been increasingly implicated in food intake control (Wang et al., 2025), particularly reward-motivated behavior (Ge and Balleine, 2022; Huijgens et al., 2024), such as impulsive operant responding (Kim et al., 2018). As a 2nd-order target site in these pathways, the BST may serve as a point of convergence for hippocampal outputs related to motivational context and internal state. The confluence of CA1v-derived neural activity on the BST may shape downstream processing within energetic- and reward-related circuits.

Previous work detailing the connectivity of CA1v neurons suggests that the majority of these cells are single projection neurons (Gergues et al., 2020). For example, only ∼25% of CA1v neurons project to two or more downstream targets, with only ∼3% of PFC projecting neurons also projecting to the LHA, ∼6% of PFC neurons also projecting to the ACB, and ∼4% of ACB neurons also projecting to the LHA. While relatively few CA1v neurons are projecting to more than one region studied here, the fact that these pathways share 2nd-order projections may allow functionally distinct hippocampal outputs to converge and be integrated downstream. Gergues et al. also found an overrepresentation of CA1v neurons that project to the LS that also send projections to the mPFC or ACB. The role of the LS as a shared 2nd-order projection of CA1v-mPFC and CA1v-ACBsh pathways suggests that hippocampal input to the LS is distributed across multiple output streams and may represent an important downstream node within hippocampal circuits relevant to reward-motivated behavior.

Further, mPFC-projecting CA1v neurons have a unique transcriptional profile, specifically enriched for genes involved in metabolic processes (Gergues et al., 2020). This suggests that the CA1v-mPFC may be uniquely positioned to integrate metabolic signals with cognitive processes.

The CA1v itself is a common 2nd-order projection target in both the CA1v-mPFC and CA1v-LHA pathways, thus providing synaptic-driven evidence of a recurrent circuit connection. Furthermore, we know from previous studies that the CA1v neurons which project to the ACBsh themselves receive projections from the LHA (Noble et al., 2019), and results above show that the CA1v-ACBsh pathway sends 2nd-order projections to the LHA as well. Thus, the CA1v is a shared downstream region of interest in all three CA1v-derived pathways. Previous research on recurrent circuits have identified feedback loops based on structure alone. Advancing these findings, the novel ATLAS approach allows us to identify recurrent CA1v circuitry based on physiological synaptic communication. To our knowledge, this is the first study identifying definitive recurrent circuit neurons driven by empirical results vs. inference made from common/shared known connections. This recurrent connectivity suggests the presence of feedback loops from and to the vHPC, which may allow hippocampal activity to be dynamically modulated by downstream striatal, cortical, and hypothalamic inputs. While we do not know which specific neurons in the CA1v these pathways projection onto, it is possible that inhibitory interneurons (IIs) within the CA1v participate in these feedback loops, allowing for fine tuning of glutamatergic hippocampal output via intra-hippocampal inhibitory feedback. IIs are known to play a crucial role in regulating circuit dynamics in the HPC for memory, fine tuning both the firing timing and frequency of primary cells in the HPC (Topolnik and Tamboli, 2022). It is possible that 2nd-order downstream projections recurrently connect back to the CA1v to inhibitory interneurons to fine tune hippocampal output based on motivational or homeostatic state.

Pathway-specific transsynaptic retrograde tracing revealed significant input to the CA1v from the intrinsic hippocampal regions, with the CA3v not surprisingly being a primary source of input to CA1v in both the CA1v-mPFC and CA1v-LHA pathways. However, while the CA1v-ACBsh pathway also received some synaptic input from the CA3v, a larger portion of its inputs were cortical, coming from the posterior amygdalar nucleus (PA), cortical nucleus of the amygdala (COA), and the basolateral amygdalar nucleus (BLA). Previous research has shown a role for amygdalar inputs to the ACB in regulating reward motivated behavior (Millan et al., 2017; Poggi et al., 2024; Taniguchi et al., 2026), and one study found that long-term potentiation (LTP) at the vHPC-ACB synapse was contingent to BLA-ACB activation (Yu et al., 2022). In this case, it was concluded that vHPC inputs to the ACB acquire increase circuit representation via the contingent BLA-ACB activation. While our identified pathway is not a direct amygdala to ACB circuit, perhaps these amygdalar inputs to the CA1v-ACBsh pathway integrate emotion and motivational context with recurrent hippocampal output pathways to influence accumbal processing. Additionally, the presence of midbrain inputs, such as the VTA in the CA1v-mPFC pathway, suggests that internal state signals related to motivation may modulate hippocampal-prefrontal communication.

There are limitations in the methodology of the current study worth noting. In the transsynaptic anterograde approach, the quantification of axon density ratings is inherently qualitative and pathway specific, with axon density within a pathway rated on a scale of 1 to 7 relative to input within (but not between) that pathway. This makes it impossible to directly compare measures of axon density between pathways, as the most and least dense projection targets are relative to each pathway, serving as a limitation in our anterograde tracing quantitative approach. Additionally, while dense axonal labelling likely indicates some terminal field presence, our approach did not resolve terminals from axons of passage (e.g., via co-labeling of presynaptic terminal markers). For the transsynaptic retrograde tracing, the current viral technology does not allow us to distinguish targeted neurons within the CA1v, leaving open the possibility that there are intra-CA1v projections – i.e. distinct CA1v neurons which give input to our primary CA1v neurons – in these pathways that we cannot identify from this approach. Additionally, neither the rabies virus nor the retro-cre-inducing virus used in our retrograde tracing approach are mediated by endogenous synaptic transmission.

In summary, our results reveal downstream and upstream neuronal projections from three distinct, multi-node pathways anchored in the CA1v. The CA1v-mPFC, CA1v-ACBsh, and CA1v-LHA pathways share common upstream targets, namely the CA3v and SUBv, though the CA1v-ACBsh pathway receives substantial innervation from cortical areas. Additionally, these projection pathways share common downstream targets, namely the LS for the CA1v-mPFC and CA1v-ACBsh projections, and the BST for the CA1v-ACBsh and CA1v-LHA projections. These common downstream targets suggest that anatomically distinct CA1v-centered pathways converge onto shared downstream nodes, potentially enabling distributed yet coordinated influence over downstream subcortical circuits involved in motivation and internal state regulation. Finally, the CA1v itself is a shared downstream projection site of the CA1v-mPFC and CA1v-LHA pathways, and such recurrent/bidirectional connectivity suggests the existence of vHPC recurrent circuits that allow CA1v activity to be reciprocally and dynamically modulated by downstream inputs. Future research should investigate how environmental exposures might affect the connections and strength of connections within these pathways, particularly exposure to a Western diet (WD). Previous research suggests that early development, and the hippocampus specifically, are particularly sensitive to WD exposure (Tsan et al., 2022; Hayes et al., 2024b, 2024a) and future research should examine if and how early-life WD exposure effects development of these motivational control-relevant connectivity circuits. Additionally, future studies should aim to genetically phenotype first-order projection neuron targets in these pathways, allowing us to better understand the nature of the recurrent connectivity discovered in some of these pathways.

---

**Supplemental Figure 1.**
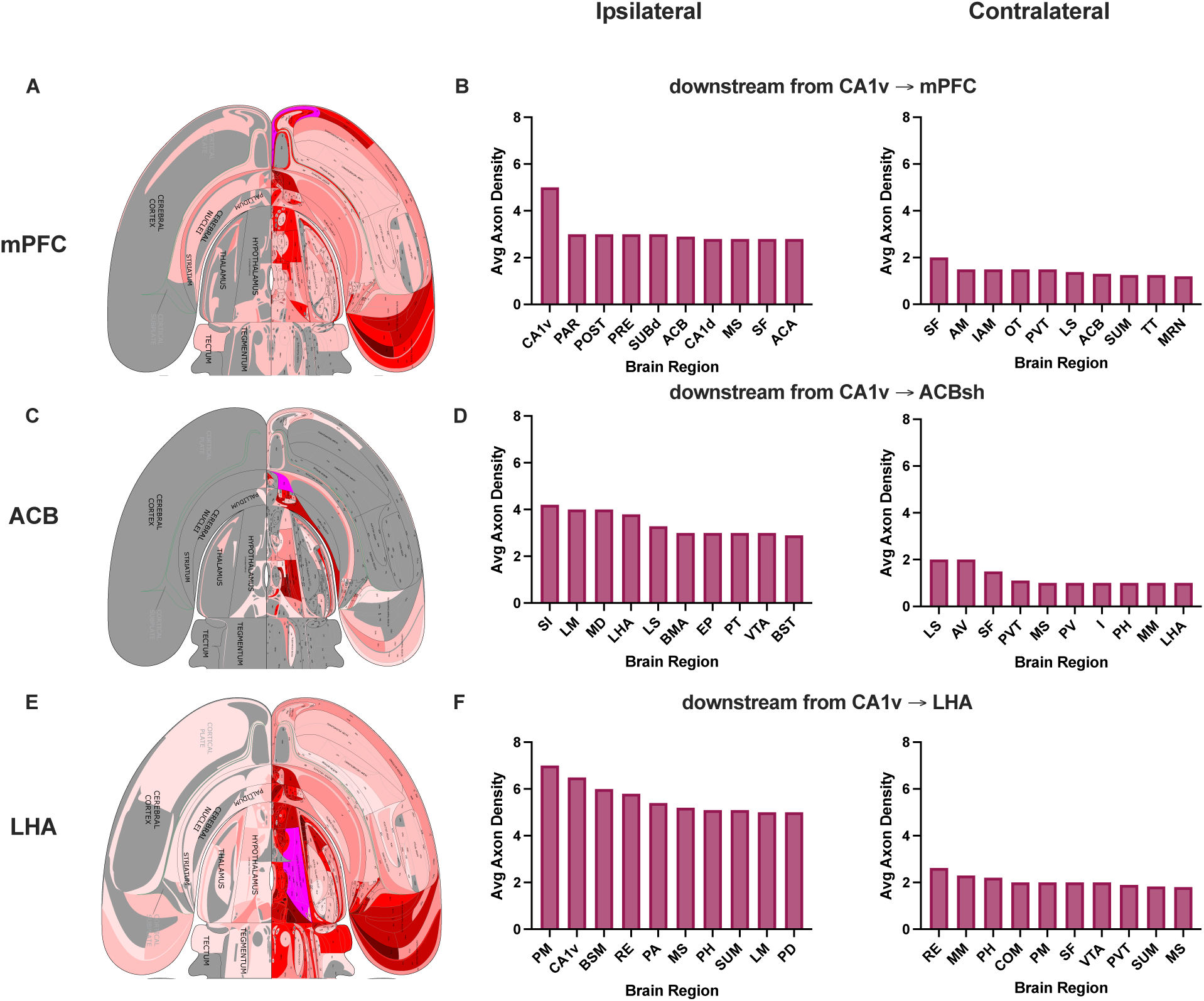
Average axon density for anterograde tracing of 2nd-order projections from CA1v pathways. **A.** Flatmap depicting average axon density for CA1v-mPFC anterograde tracing. **B**. Ipsilateral (left) and contralateral (right) graphs depicting average axon density rating in CA1v-mPFC pathway. **C.** Flatmap depicting average axon density for CA1v-ACB anterograde tracing. **D**. Ipsilateral (left) and contralateral (right) graphs depicting average axon density rating in CA1v-ACB pathway. **E**. Flatmap depicting average axon density for CA1v-LHA anterograde tracing. **F**. Ipsilateral (left) and contralateral (right) graphs depicting average axon density rating in CA1v-LHA pathway.

**Supplemental Figure 2.**
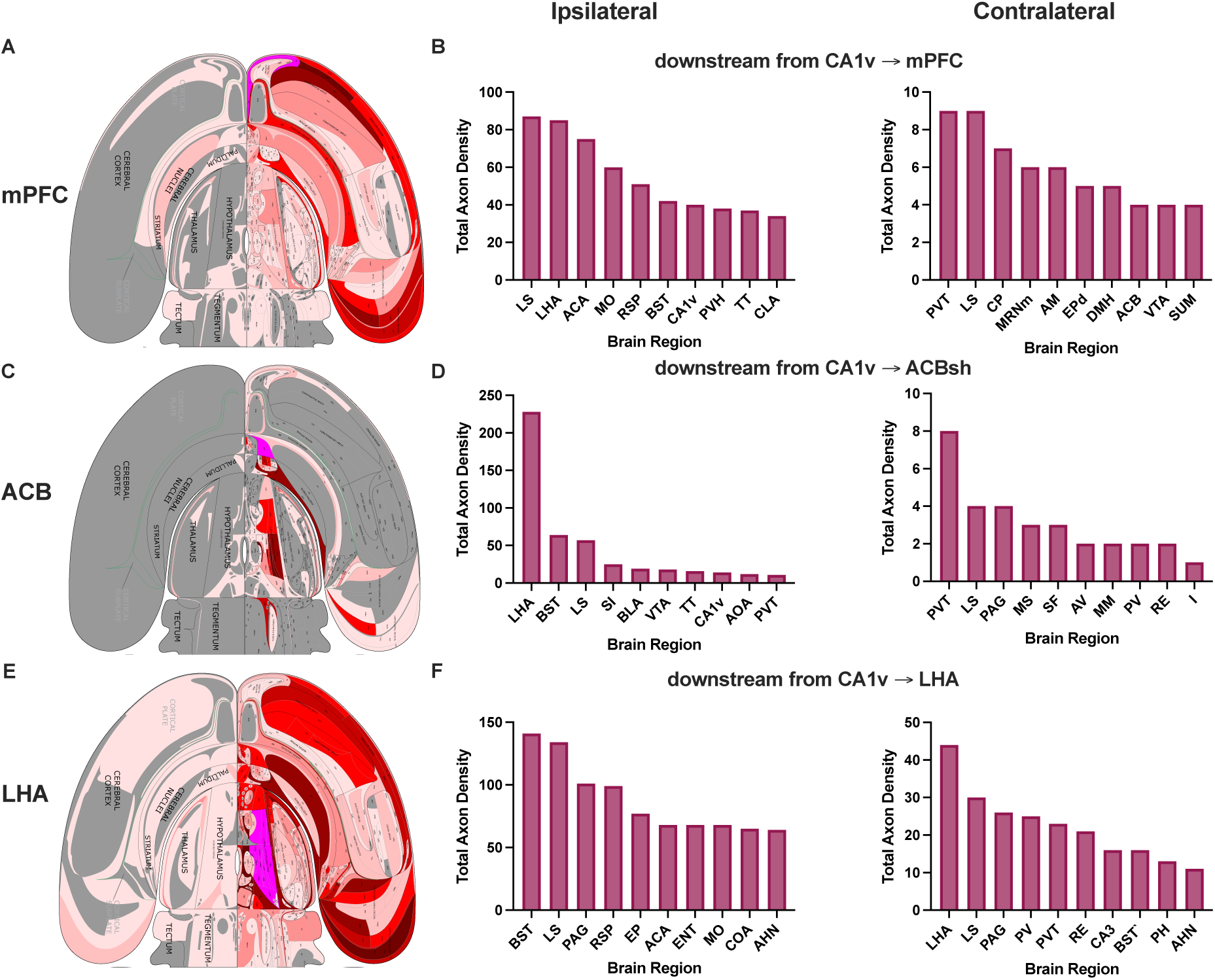
Total axon density for anterograde tracing of 2nd-order projections from CA1v pathways. **A.** Flatmap depicting total axon density for CA1v-mPFC anterograde tracing. **B**. Ipsilateral (left) and contralateral (right) graphs depicting total axon density rating in CA1v-mPFC pathway. **C.** Flatmap depicting total axon density for CA1v-ACB anterograde tracing. **D**. Ipsilateral (left) and contralateral (right) graphs depicting total axon density rating in CA1v-ACB pathway. **E**. Flatmap depicting total axon density for CA1v-LHA anterograde tracing. **B**. Ipsilateral (left) and contralateral (right) graphs depicting total axon density rating in CA1v-LHA pathway.

**Supplemental Figure 3.**
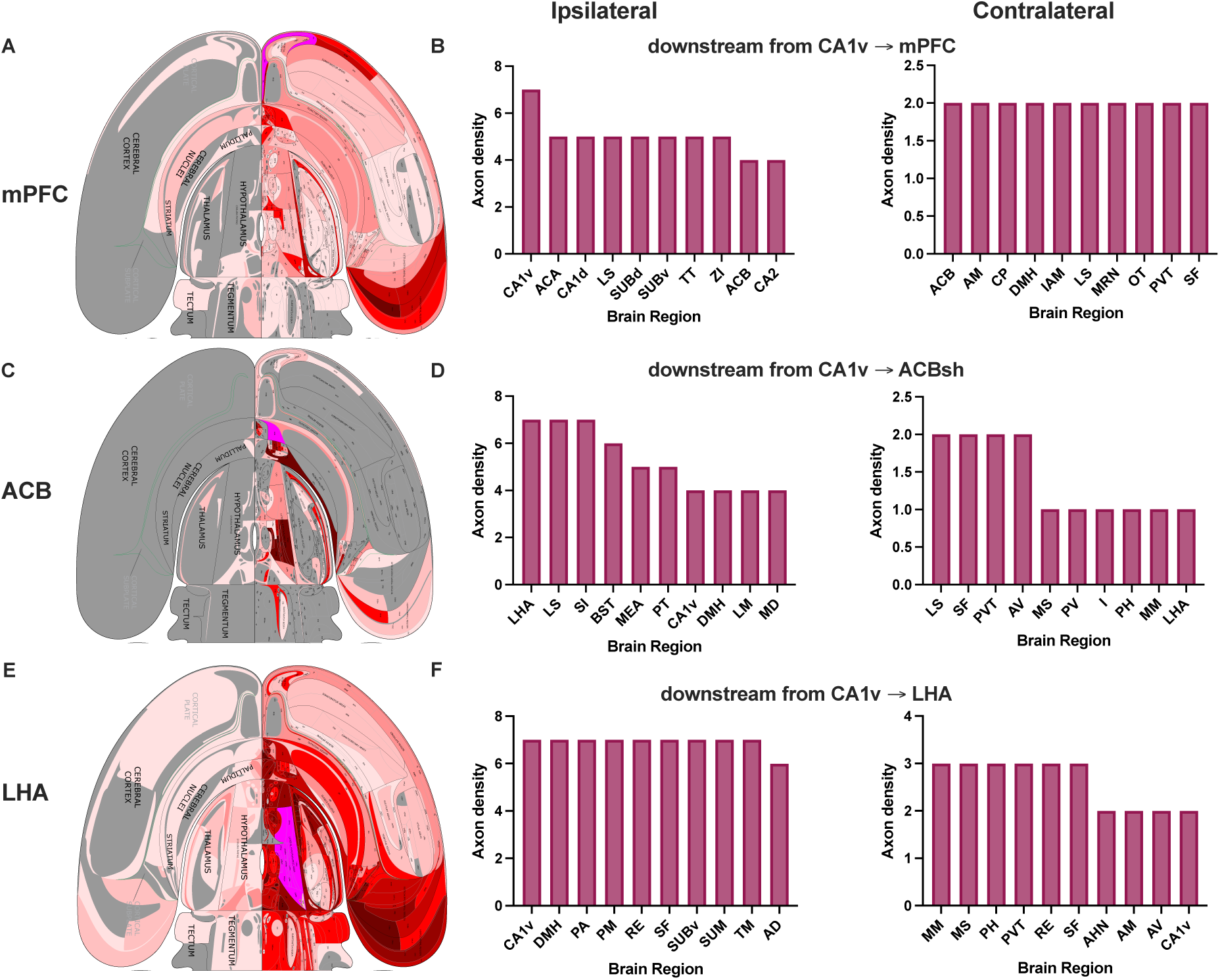
Maximum axon density for anterograde tracing of 2nd-order projections from CA1v pathways. **A.** Flatmap depicting maximum axon density for CA1v-mPFC anterograde tracing. **B**. Ipsilateral (left) and contralateral (right) graphs depicting maximum axon density rating in CA1v-mPFC pathway. **C.** Flatmap depicting maximum axon density for CA1v-ACB anterograde tracing. **D**. Ipsilateral (left) and contralateral (right) graphs depicting maximum axon density rating in CA1v-ACB pathway. **E**. Flatmap depicting maximum axon density for CA1v-LHA anterograde tracing. **B**. Ipsilateral (left) and contralateral (right) graphs depicting maximum axon density rating in CA1v-LHA pathway.

**Supplemental Figure 4.**
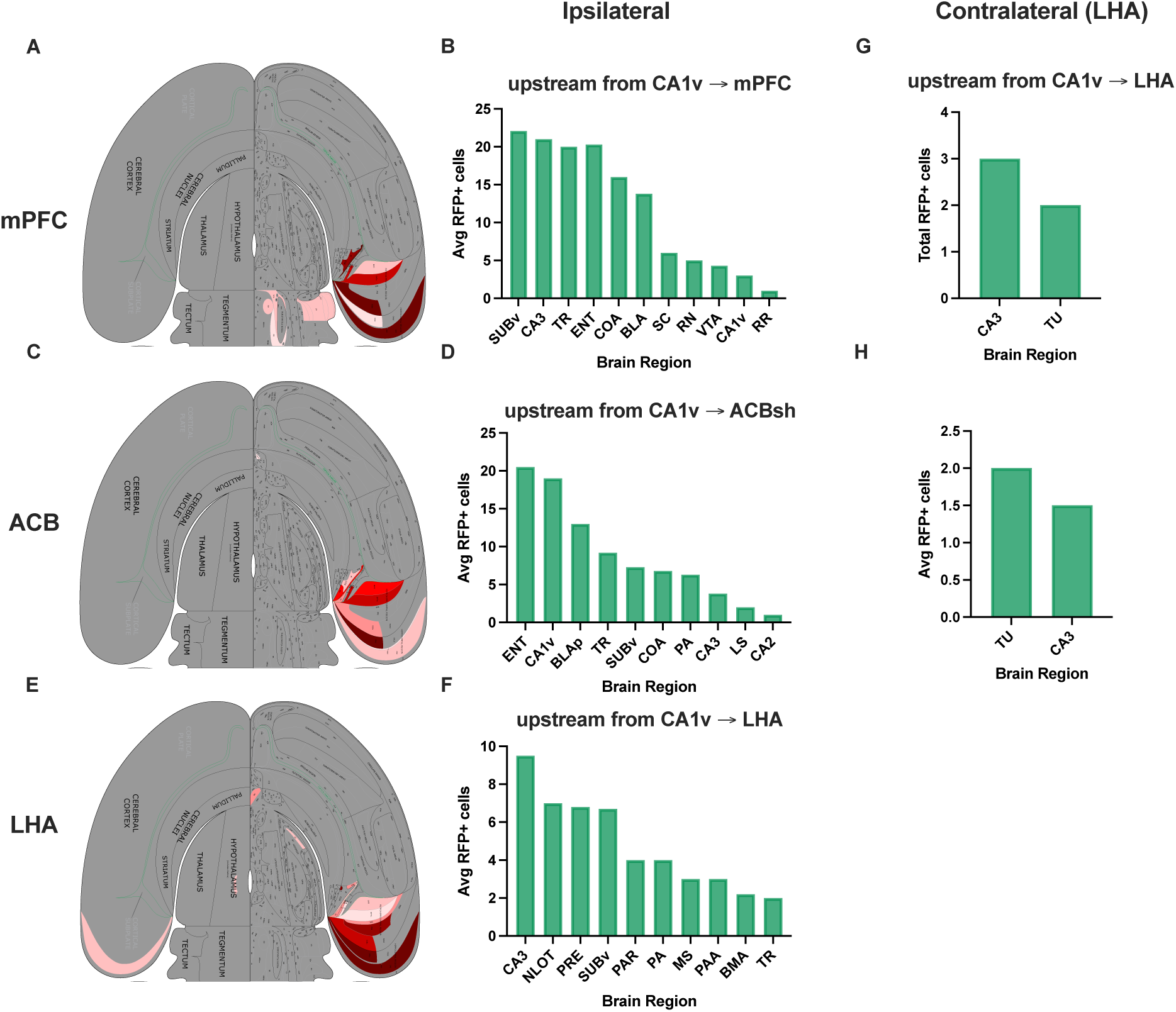
Average RFP+ cell bodies in retrograde tracing of primary inputs to CA1v pathways. **A.** Flatmap depicting average RFP+ cell bodies for CA1v-mPFC retrograde tracing. **B**. Ipsilateral graph depicting average RFP+ cells for retrograde tracing in CA1v-mPFC pathway. **C.** Flatmap depicting average RFP+ cell bodies for CA1v-ACB retroograde tracing. **D**. Ipsilateral graph depicting average RFP+ cells for retrograde tracing in CA1v-ACB pathway. **E.** Flatmap depicting average RFP+ cell bodies for CA1v-LHA retrograde tracing. **F**. Ipsilateral graph depicting average RFP+ cells for retrograde tracing in CA1v-LHA pathway. **G.** Contralateral graph depicting total RFP+ cells for retrograde tracing in CA1v-LHA pathway. **H.** Contralateral graph depicting total RFP+ cells for retrograde tracing in CA1v-LHA pathway.

